# Sequence-dependent transferability of the LRLLR membrane translocation motif: A computational study of smacN and NR2B9c peptides

**DOI:** 10.64898/2026.02.19.706823

**Authors:** D. Muñoz-Gacitúa, Jenny Blamey

## Abstract

The LRLLR cell-penetrating motif can be transferred to confer membrane translocation activity, but only to compatible recipient peptides. Using umbrella sampling molecular dynamics simulations, we demonstrate that C-terminal LRLLR addition to the pro-apoptotic smacN peptide eliminates its translocation barrier entirely, transforming a +65 kJ/mol barrier into a −50 kJ/mol energy well. In contrast, N-terminal LRLLR addition to the neuroprotective NR2B9c peptide increases the translocation barrier from +85 to +100 kJ/mol, demonstrating that motif transfer can prove counterproductive for incompatible sequences.

Cell-penetrating peptides offer promising strategies for intracellular delivery of therapeutic cargo, yet the sequence determinants governing their activity remain incompletely understood. The LRLLR motif, identified through systematic screening as essential for spontaneous membrane translocation, represents a minimal penetrating element whose transferability has not been previously evaluated. We appended this motif to two clinically relevant peptides: smacN, a tetrapeptide targeting inhibitor of apoptosis proteins in chemotherapy-resistant cancers, and NR2B9c, a nonapeptide that disrupts excitotoxic signaling in ischemic stroke.

Potential of mean force profiles calculated across a POPC/POPG bilayer, combined with analysis of hydrogen bonding patterns, secondary structure propensity, and conformational dynamics, reveal the structural basis for these divergent outcomes. Successful transfer to smacN results from favorable complementarity: the hydrophobic, neutral smacN provides an ideal platform for the charged, amphipathic LRLLR motif, yielding a chimera capable of simultaneous interaction with both membrane leaflets. Transfer failure with NR2B9c stems from conformational rigidity induced by intramolecular hydrogen bonding, which prevents optimal membrane insertion, combined with unfavorable positioning of internal polar residues at the bilayer center.

These findings establish that cell-penetrating motif transfer requires compatibility in charge distribution, hydrophobicity, and conformational flexibility between the motif and recipient sequence. The smacN-LRLLR chimera emerges as a promising candidate for experimental validation as a membrane-permeable therapeutic for survivin-positive tumors. More broadly, this work demonstrates the value of computational screening to identify compatible motif-cargo pairings prior to experimental investment.

## Introduction

The phospholipid bilayer of the cell membrane presents a formidable barrier to polar drug candidates, effectively excluding peptide therapeutics and macromolecules that lack the hydrophobicity or specific transport mechanisms required for passive diffusion^1–3^. Consequently, numerous promising therapeutic peptides, proteins, and nucleic acid conjugates have failed to reach clinical application due to their inability to achieve cytosolic concentrations sufficient for therapeutic efficacy. ^4–7^.

Among these, cell-penetrating peptides (CPPs) have emerged as promising molecular vectors. These short sequences, typically comprising 5–30 amino acids, possess the unique ability to traverse plasma membranes while transporting diverse cargoes including proteins, nucleic acids, and small molecules^8,9^. First identified in 1988 with the discovery that the HIV-1 TAT protein could translocate across membranes^10^, followed by the characterization of the Antennapedia homeodomain-derived Penetratin^11,12^, CPPs established a new paradigm for intracellular delivery. Initially believed to enter cells via endocytosis, subsequent methodological advances revealed that these peptides are quite more complex in their mechanistical nature, as they usually feature more than one translocation mechanism^13^, and which one they employ is heavily multifactorial^14–16^.

Despite their translocation capability, CPPs exhibit significant limitations that hinder their clinical translation^17–19^. Their delivery efficiency is highly cargo-dependent, with large proteins and nucleic acids frequently becoming trapped in endosomal compartments rather than reaching the cytosol^20^. Furthermore, cellular uptake mechanisms remain unpredictable, varying with concentration, membrane composition, and sequence context in ways that are not yet fully understood. These constraints underscore the urgent need to identify specific sequence motifs that govern membrane translocation and determine whether such elements can be rationally transferred to confer cell-penetrating ability to therapeutic peptides.

The identification of specific sequence motifs responsible for membrane translocation has proven challenging, as CPP activity often emerges from complex interplay between charge distribution, hydrophobicity, and secondary structure propensity. However, recent systematic screening approaches have begun to isolate minimal penetrating elements^21–23^. Marks and colleagues developed a method to identify spontaneously translocating CPPs by screening randomly generated peptide libraries against protein-free membrane mimetics^24^, ensuring that positive hits must cross the bilayer without mediation by active biological processes. This approach yielded a set of spontaneously translocating CPPs, among which the peptides TP1 and TP2 demonstrated exceptional penetrating capability. Notably, analysis of successful sequences revealed a recurring motif: the pentapeptide LRLLR appeared frequently among peptides capable of crossing membranes without disrupting bilayer integrity.

Subsequent biophysical characterization revealed mechanistic details of LRLLR-mediated translocation. When TP1 and TP2 were incubated with vesicles bearing the fluorophore NBD on the outer leaflet, fluorescent lipids redistributed to the inner leaflet, indicating that peptide translocation involves a coupled lipid flip-flop mechanism^25^. These findings were confirmed and extended through nuclear magnetic resonance spectroscopy combined with molecular dynamics simulations^26^. This combined approach demonstrated that TP1 occupies a significant potential energy well when embedded in the membrane, and critically, that the spacing between the two guanidinium groups of the arginine residues is essential for penetrating activity. The authors proposed that each guanidinium moiety interacts simultaneously with opposite leaflets of the bilayer, creating a tensioned configuration that serves as a transition state facilitating lipid flip-flop and consequent translocation. The hydrophobic leucine residues intervening between the arginines are thought to stabilize this transition state, accelerating translocation by reducing exposure of the charged groups to the hydrophobic core.

While substantial evidence points to the LRLLR sequence as responsible for the penetrating activity of TP1 and TP2^22,27^, it remains unconfirmed whether this capability can be transferred to other peptides. The present study addresses this question by appending the LRLLR motif to two therapeutically promising peptides: smacN and NR2B9c. These candidates were selected not only for their clinical relevance but also because their target recognition sites permit modification at opposite termini, enabling us to test motif transfer at both C-terminal and N-terminal positions.

The smacN peptide comprises the four N-terminal residues (AVPI) of the Smac/DIABLO protein, a mitochondrial factor that promotes apoptosis by neutralizing inhibitor of apoptosis proteins (IAPs)^28^. This minimal tetrapeptide retains capacity to bind and inhibit XIAP and survivin^29,30^. Survivin overexpression occurs in numerous carcinogenic tumors, where elevated concentrations correlate with resistance to chemotherapeutic treatment^31–33^. The smacN peptide therefore represents a promising therapeutic agent for survivin-positive tumors, provided it can achieve sufficient intracellular concentrations^34^. Since smacN is recognized by its N-terminus at the IAP binding groove^35^, modifications must be introduced at the C-terminal end, making this peptide an ideal candidate to test whether C-terminal LRLLR addition confers membrane-penetrating activity.

The NR2B9c peptide comprises the nine C-terminal residues (KLSSIESDV) of the NR2B subunit of the NMDA receptor. Under excitotoxic conditions, excessive glutamate signaling through NMDA receptors triggers a pathological cascade: the scaffolding protein PSD-95 couples activated NR2B subunits to neuronal nitric oxide synthase (nNOS), leading to toxic nitric oxide production and subsequent ischemic damage^36–38^. NR2B9c competitively disrupts the PSD-95/NR2B interaction, thereby interrupting this neurotoxic signaling pathway without affecting normal NMDA receptor function^39,40^. Because target recognition occurs through the C-terminal PDZ-binding motif, modifications must be introduced at the N-terminus, making NR2B9c suitable for testing N-terminal LRLLR addition.

The therapeutic potential of NR2B9c has been evaluated clinically through conjugation with the TAT cell-penetrating peptide. The resulting construct, NA-1 (also known as nerinetide), underwent Phase III clinical trials for acute ischemic stroke treatment^41,42^. Unfortunately, the ESCAPE-NA1 trial failed to demonstrate that NA-1 reduced infarct volume compared to a placebo, with evidence that alteplase (a thrombolytic drug delivered to all patients in the trial) was inhibiting the effect of the drug being tested. A subsequent trial, ESCAPE-NEXT, is investigating NA-1 in patients treated with endovascular thrombectomy alone^43^. These clinical results establish proof-of-concept for NR2B9c efficacy but highlight the continued need for optimized delivery strategies. Whether the shorter LRLLR motif can provide sufficient membrane translocation enhancement—potentially with reduced immunogenicity and improved pharmacokinetics compared to the longer TAT sequence—remains an open question.

To evaluate the translocation-enhancing potential of the LRLLR motif, we employed umbrella sampling molecular dynamics simulations to calculate the potential of mean force (PMF) for peptide translocation across a model phospholipid bilayer. This computational approach applies a series of harmonic restraints to position the peptide at defined distances from the bilayer center, sampling conformational space at each position before reconstructing the complete free energy profile using the weighted histogram analysis method (WHAM). Umbrella sampling has been extensively validated for membrane translocation studies, successfully reproducing experimental translocation rates and partition coefficients for amino acid side chain analogs^44^, antimicrobial peptides^45^, and cell-penetrating peptides including oligoarginines and penetratin^46^.

The computational approach offers several advantages for the present investigation. First, PMF calculations provide quantitative energetic information—barrier heights and well depths—that can be directly related to translocation feasibility and membrane partitioning. Second, the atomistic detail of molecular dynamics simulations reveals structural mechanisms underlying energetic profiles, including hydrogen bonding patterns, secondary structure changes, and membrane perturbations that would be difficult to access experimentally. Third, computational screening enables direct comparison of modified and unmodified peptides under identical conditions, isolating the contribution of the LRLLR motif without confounding variables introduced by different experimental preparations or measurement techniques.

This study aims to determine whether the LRLLR motif functions as a transferable translocation module capable of conferring membrane-penetrating activity to therapeutic peptides. We test three specific hypotheses: first, that LRLLR addition will reduce the free energy barrier for membrane translocation of both smacN and NR2B9c; second, that motif addition will preserve the structural features required for target recognition by these therapeutic peptides; and third, that PMF profiles combined with structural analysis will reveal the molecular mechanisms by which LRLLR facilitates or fails to facilitate translocation. The results provide fundamental insight into the sequence determinants of CPP activity and offer practical guidance for the rational design of cell-penetrating therapeutic peptide conjugates.

## Methods

To evaluate the translocation-enhancing potential of the LRLLR motif, we employed a computational workflow combining steered molecular dynamics with umbrella sampling to calculate potential of mean force profiles for peptide translocation across a model phospholipid bilayer. The workflow proceeded in three stages: first, peptide and membrane systems were prepared and equilibrated independently to obtain stable initial configurations; second, steered molecular dynamics simulations generated translocation pathways by pulling each peptide through the bilayer at constant velocity; third, configurations extracted along these pathways served as starting points for umbrella sampling simulations, from which PMF profiles were reconstructed using the weighted histogram analysis method. All simulations employed the CHARMM36 force field and were performed using GROMACS 2024. Structural and energetic analyses, including hydrogen bonding patterns, secondary structure propensity, and conformational fluctuations, were extracted from the umbrella sampling trajectories to characterize the molecular mechanisms underlying the calculated free energy profiles.

### Molecular dynamics parameters

Molecular dynamics simulations were carried out using GROMACS in its 2024 version^47^. CHARMM36 forcefield was chosen as it has been benchmarked against a wide array of experimental data for both lipids and peptides, showing accurate reproduction of lipid phase behavior and peptide secondary-structure propensities^48^. Periodic box conditions were applied to the three dimensions of the simulation. A modified Berendsen thermostat was used to maintain the temperature to 310 K during the simulation, and a semi-isotropic Parrinello-Rahman barostat was used to maintain the pressure to 1 bar. During equilibration simulations, a 1 fs time-step was used for integration to lower the chance of an *exploding* trajectory as they begin from a configuration that is far from equilibrium. After this simulation has relaxed the configuration to a state with proper contacts, density and pressure; a production simulation is performed with a 2 fs time-step. On these production simulations, a LINCS constraint algorithm is applied to bonds containing hydrogen atoms to maintain correct bond lengths^49^. Van-der-waals and coulombic interactions were calculated fully between particles closer than 1.2 nm. Long range interactions were calculated using a Particle-mesh ewald (PME) summation.

### System composition and structure preparation

The initial structure of each peptide was built using Bio2Byte PeptideBuilder library^50^, the exact sequence of each peptide is specified in table 1. Each peptide initial structure was submitted to a 100 ns simulation in water to allow them to acquire a natural and stable secondary structure, which was verified through RMSD measurements.

**Table 1:** Peptides used in this study and their sequences.

| **Peptide** | **Sequence** |
| --- | --- |
| *smacN* | AVPI |
| *smacN-LRLLR* | AVPILRLLR |
| *NR2B9c* | KLSSIESDV |
| *LRLLR-NR2B9c* | LRLLRKLSSIESDV |

The membrane was simulated with a 90% POPC and 10% POPG symmetrical composition. [rationale for this proportion]. The proportion was achieved placing 8 POPG and 72 POPC molecules in a bilayer configuration of 5 nm by 5 nm. Initial configuration of the membrane was built using Packmol^51^. The simulation box was then enlarged in the Z axis, perpendicular to the bilayer, up to 9nm and then filled with water and ion molecules. Na+ and Cl-ions were added to neutralize the charges of the lipids, and 0.1M worth of ions on top of that. The size of the box was chosen so peptides in the simulation could not interact with a copy of itself in the periodic box conditions. Finally, the membrane simulation box is submitted to an energy minimization procedure, followed by a 1 ns equilibration simulation and a 100 ns full simulation that allows the bilayer to reach a natural state. This was verified by checking if the thickness and area per lipid reached a stable value.

### Steered MD and umbrella sampling

Peptide translocation was simulated by pushing the peptide through the membrane in simulation. In each case, the peptide was initially placed in the aqueous portion of the simulation. Then, in a steered simulation, the peptide was pushed along the Z-axis perpendicular to the membrane at a rate of 0.2 nm/ns with a force constant of 800 kJ mol-1 nm-2. This simulation was performed until the peptide looped though the entire periodic box. From this trajectory, snapshots were taken every 0.2 nm of peptide travel, which were used as starting points for an Umbrella Sampling simulation. Each of these simulation windows was equilibrated during 1 ns, and then run though a 20 ns transient simulation before a 40 ns production simulation. On these simulations, an external force restrained the center-of-mass of the peptide along the Z-axis with a force constant of 800 kJ mol-1 nm-2. After completion of each simulation window, an automated routine checked if there were less than 3000 data points per peptide position (which translates to 300 ps worth of data), and extended the corresponding simulation if that was the case. Total simulation time per peptide is specified in table S1.

### Free energy calculation and trajectory analysis

Potential of mean force (PMF) profiles were calculated from the umbrella sampling simulations using the weighted histogram analysis method (WHAM)^52^. The GROMACS implementation (gmx wham) was estimated using the trajectory-based bootstrap method with 100 bootstrap iterations. This approach samples complete trajectories with replacement, preserving correlations within individual umbrella windows employed for all PMF reconstructions. The analysis was performed at 310 K, consistent with the simulation temperature. Statistical uncertainties were assessed sampling variability across windows^53^. Sufficient overlap between adjacent umbrella windows was confirmed for all peptide systems (Figure S2)

## Results

To investigate whether the membrane-translocating properties of the LRLLR motif can be transferred to therapeutic peptides, we performed umbrella sampling molecular dynamics simulations on three systems: the LRLLR motif alone, the smacN peptide (AVPI) with and without C-terminal LRLLR addition, and the NR2B9c peptide (KLSSIESDV) with and without N-terminal LRLLR addition. For each system, we calculated the potential of mean force (PMF) profile across a POPC/POPG (90:10) bilayer, analyzed hydrogen bonding patterns with lipids, water, and intramolecular partners, and characterized secondary structure propensity as a function of membrane depth. These analyses reveal that motif transfer success depends critically on the physicochemical properties and structural behavior of the recipient peptide.

### LRLLR motif

The isolated LRLLR pentapeptide exhibits a characteristic PMF profile with a pronounced energy well of approximately −40 kJ/mol at the membrane-water interface, located 2 nm from the bilayer center (Figure 1). Translocation requires surmounting a substantial barrier of +60 kJ/mol relative to bulk water, or approximately +100 kJ/mol relative to the interfacial minimum.

**Figure 1:**
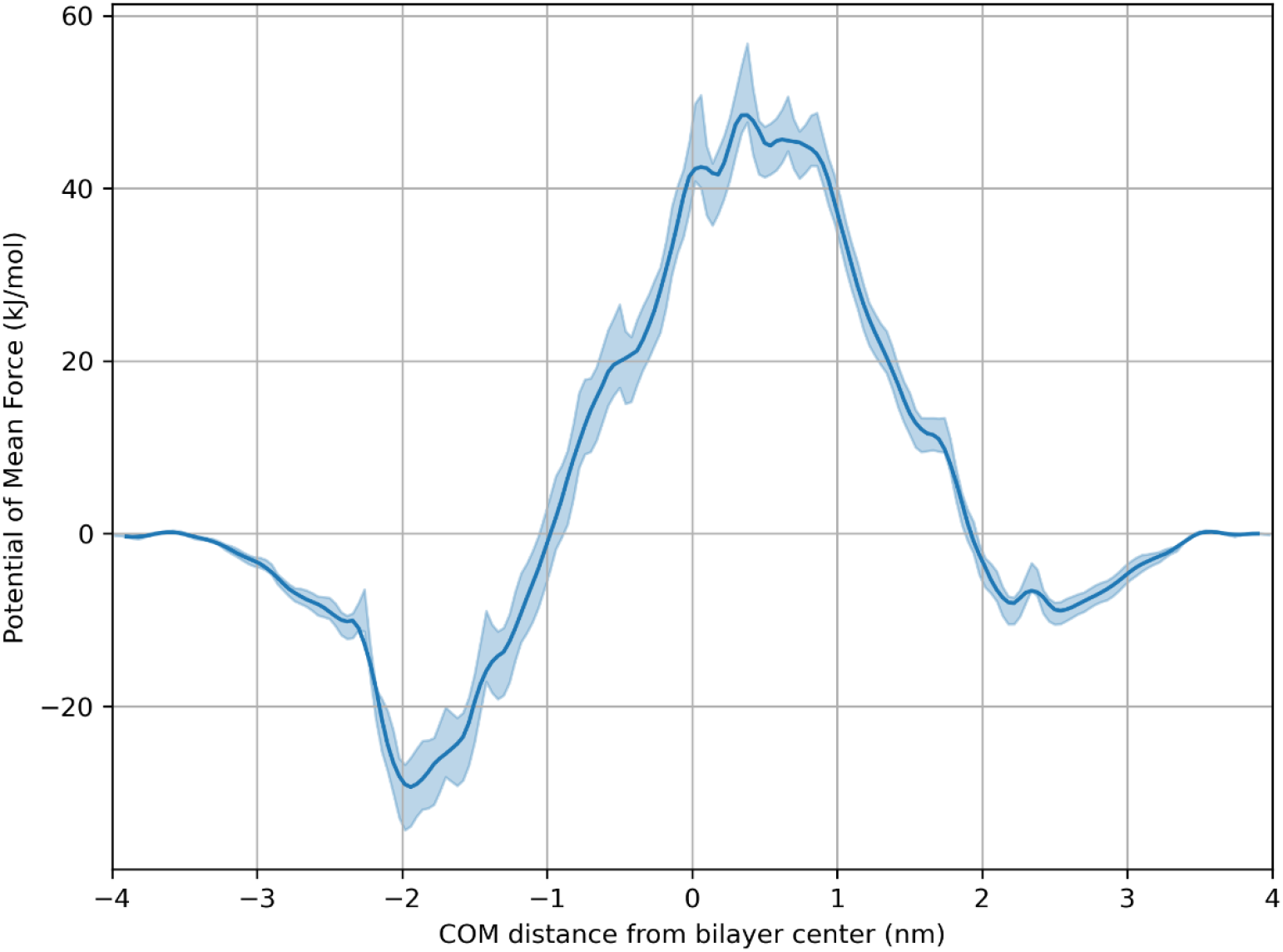
Potential of mean force profile for the LRLLR translocating a 90:10 POPC/POPG membrane. Shaded area around the profile represents a single standard deviation from the potential at that point.

At the interfacial energy minimum, the peptide adopts a relatively stable configuration as evidenced by low RMSD and RMSF values (Figure 2). Hydrogen bond analysis reveals that both arginine residues (Arg2 and Arg5) engage simultaneously with lipid headgroups and interfacial water molecules, with water hydrogen bonds predominating at this position. The peptide exhibits modest helical character, spending approximately 20% of simulation time in helical conformations at the interface—an increase from the 10% observed in bulk solution—though this propensity decreases to only 3% when the peptide is positioned within the bilayer core.

**Figure 2:**
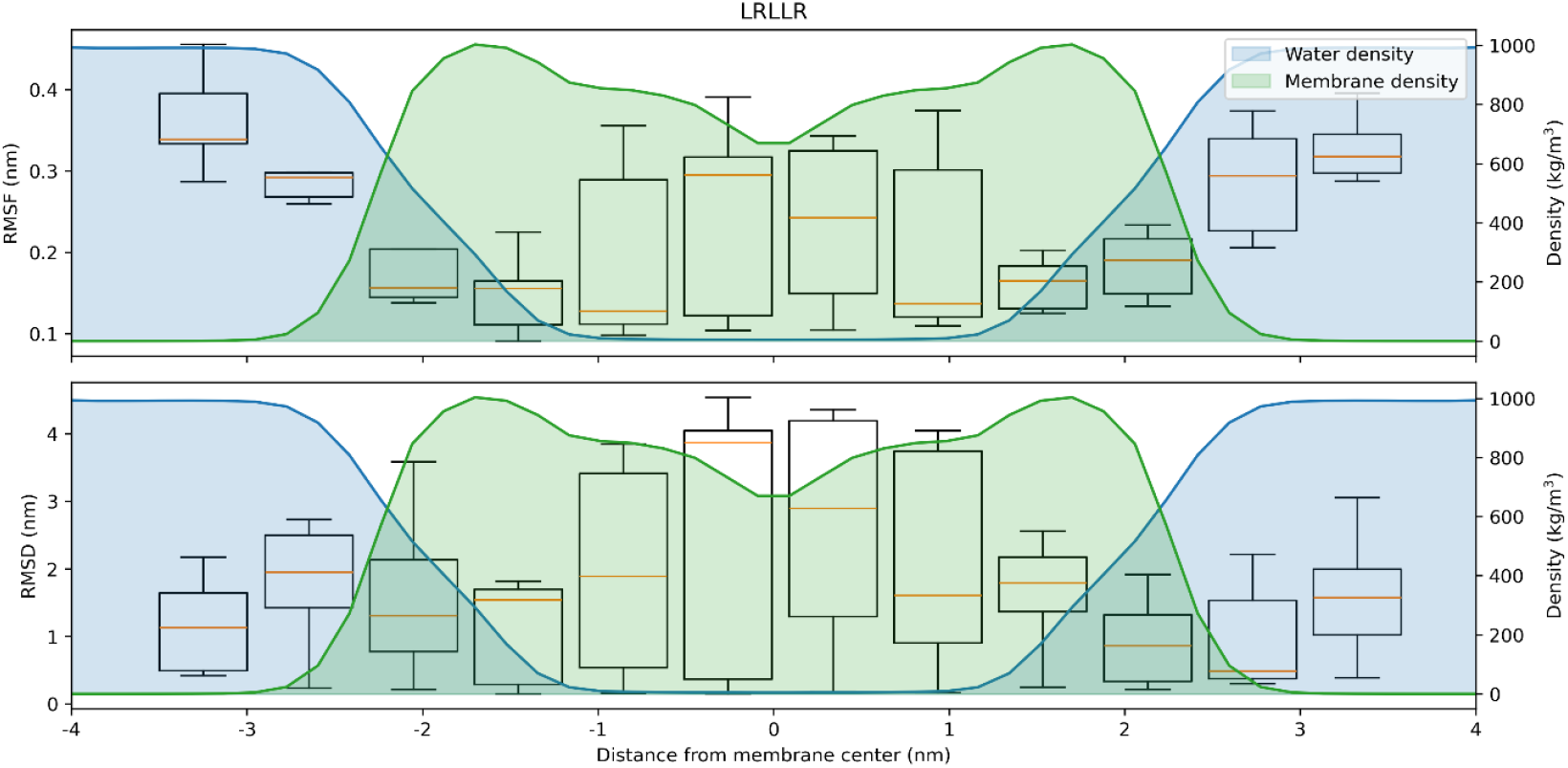
Position fluctuation plots of LRLLR peptide as a function of its distance from the bilayer center as it translocates said bilayer. Top plot shows RMSF and bottom plot shows RMSD. Shaded area in the background shows the water and lipid density as a function of the distance from the bilayer center.

**Figure 3:**
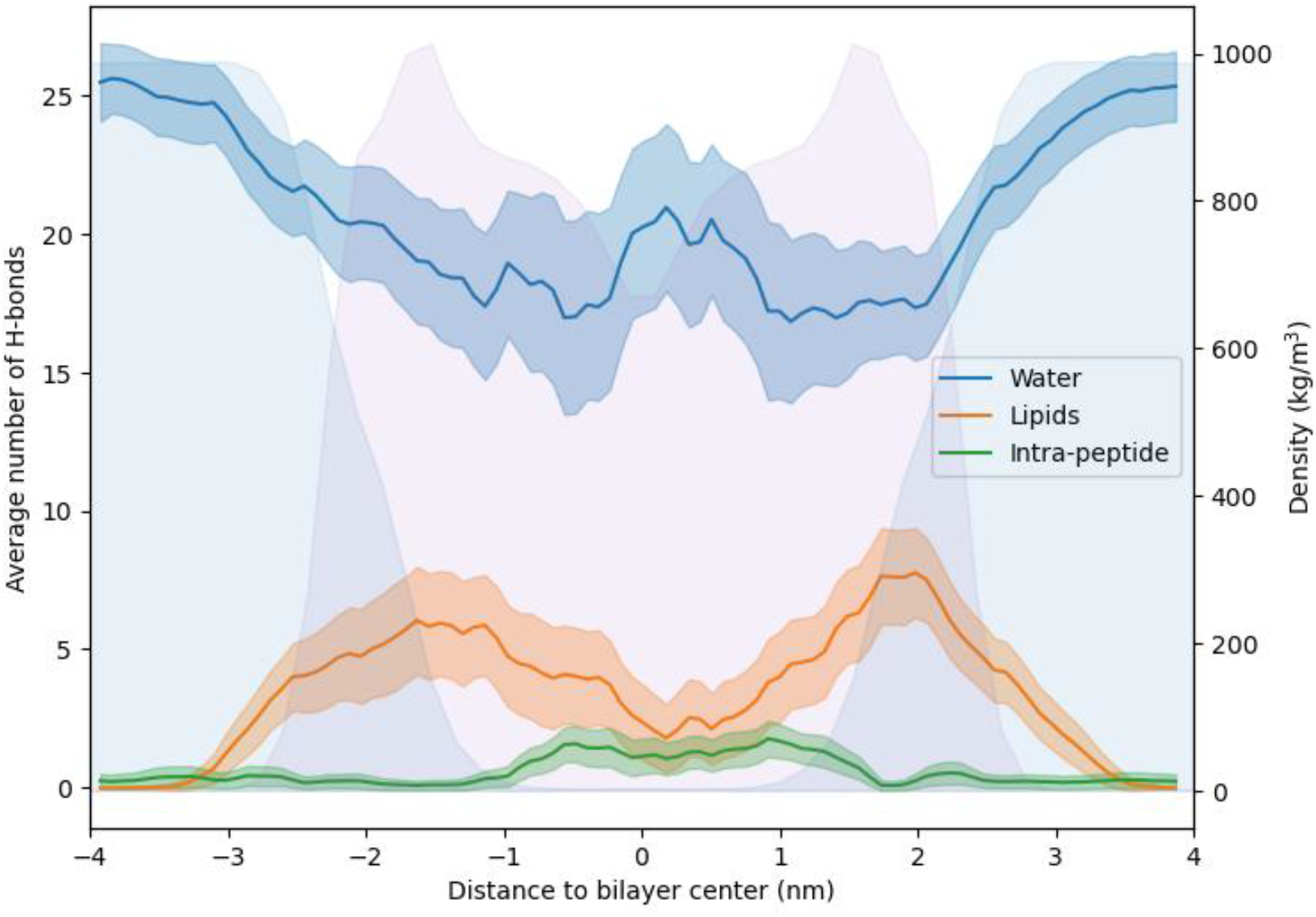
Number of hydrogen bonds involving the LRLLR peptide, separated by target. Shaded area in the background shows the water and lipid density as a function of the distance from the bilayer center.

The translocation mechanism involves passage through a transition state located approximately 0.5 nm beyond the bilayer center. At this position, both arginines form intramolecular hydrogen bonds; however, this interaction alone proves insufficient for stabilization. A transient water pore forms around the peptide, enabling 20 ± 5 hydrogen bonds with water molecules to partially stabilize this high-energy configuration. Final translocation is achieved as Leu1 and Arg2 engage with phosphate groups of the inner leaflet, effectively pulling the remainder of the peptide across the bilayer, with 7 ± 3 lipid headgroup hydrogen bonds observed during this stage.

### smacN peptide

The smacN tetrapeptide (AVPI) is a neutral, highly hydrophobic sequence (GRAVY index: 2.14) that targets inhibitor of apoptosis proteins through its N-terminus. Unmodified smacN exhibits a shallow interfacial energy well of −10 kJ/mol and faces a translocation barrier of +65 kJ/mol (Figure 4). Owing to its short length, this peptide does not adopt any discernible secondary structure throughout the translocation process.

**Figure 4:**
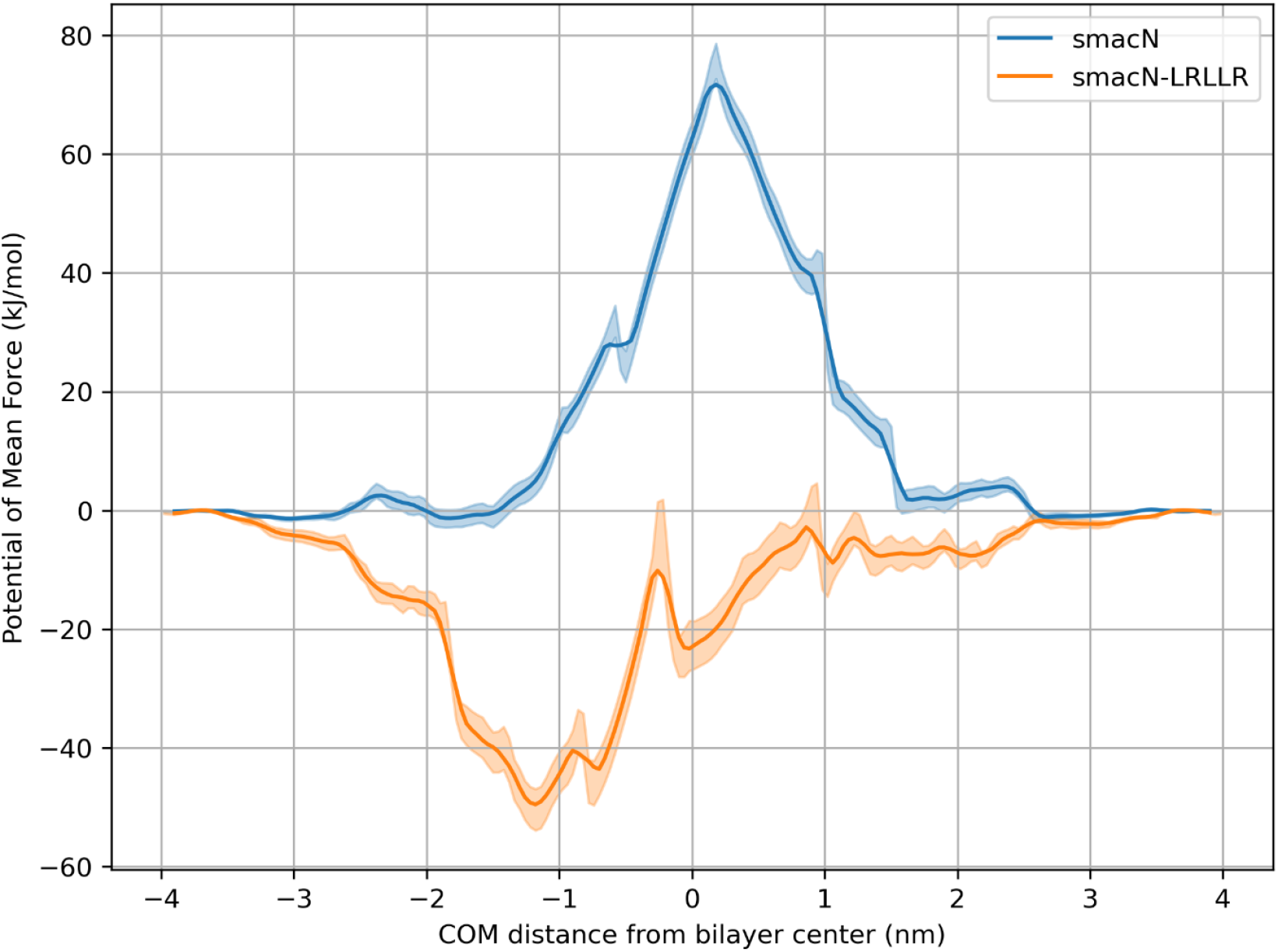
Potential of mean force profile for the smacN peptide and its chimeric variation translocating a 90:10 POPC/POPG membrane. Shaded area around the profile represents a single standard deviation from the potential at that point.

**Figure 5:**
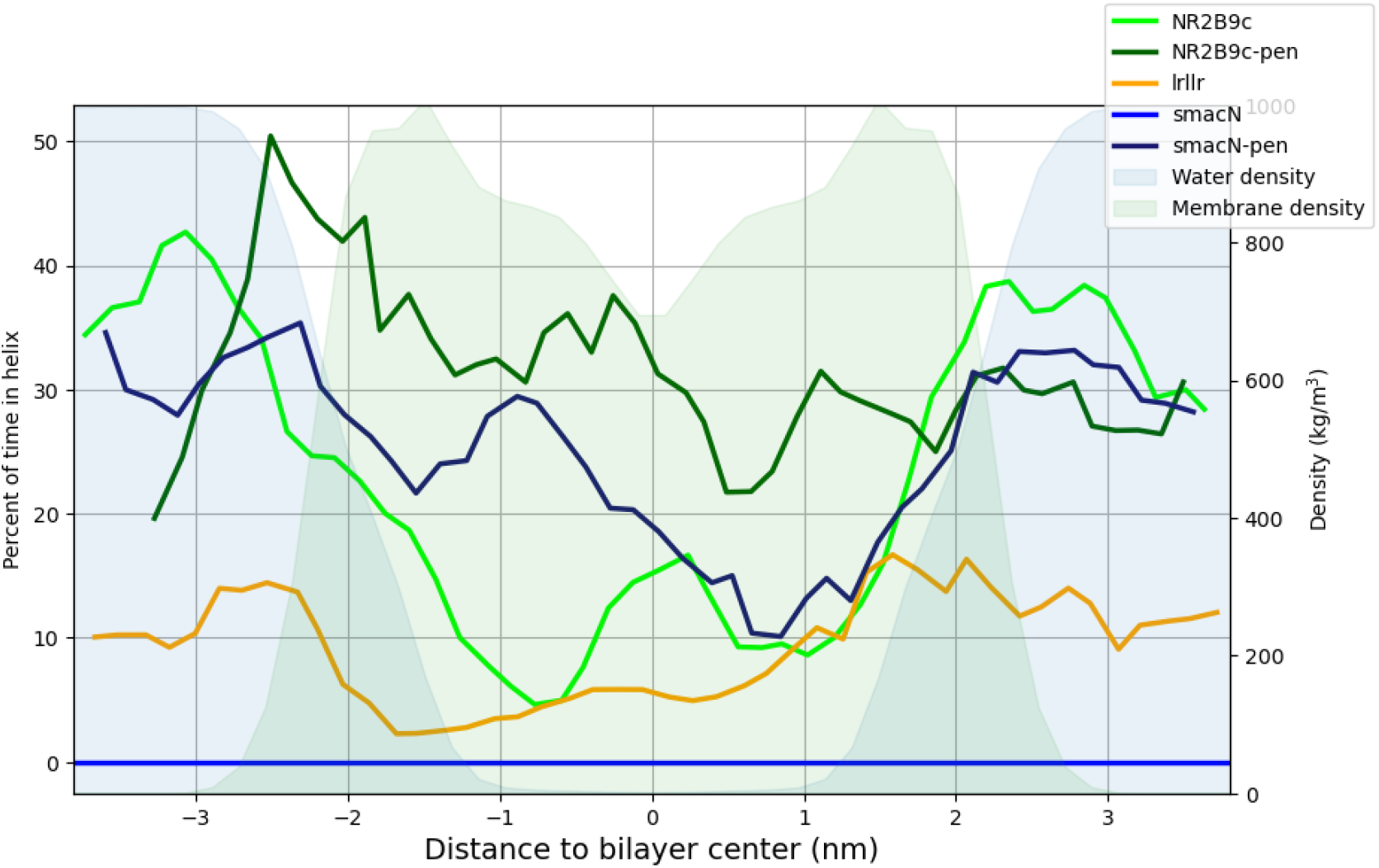
Helical propensity as a function of bilayer depth. The lines represent the fraction of simulation time spent in an α-helical conformation, determined by DSSP analysis, for each peptide. Due to the short length of the sequences, these values reflect the formation of transient helical structures rather than stable secondary folding. The shaded background illustrates the density profiles of water and lipid components relative to the bilayer center.

**Figure 6:**
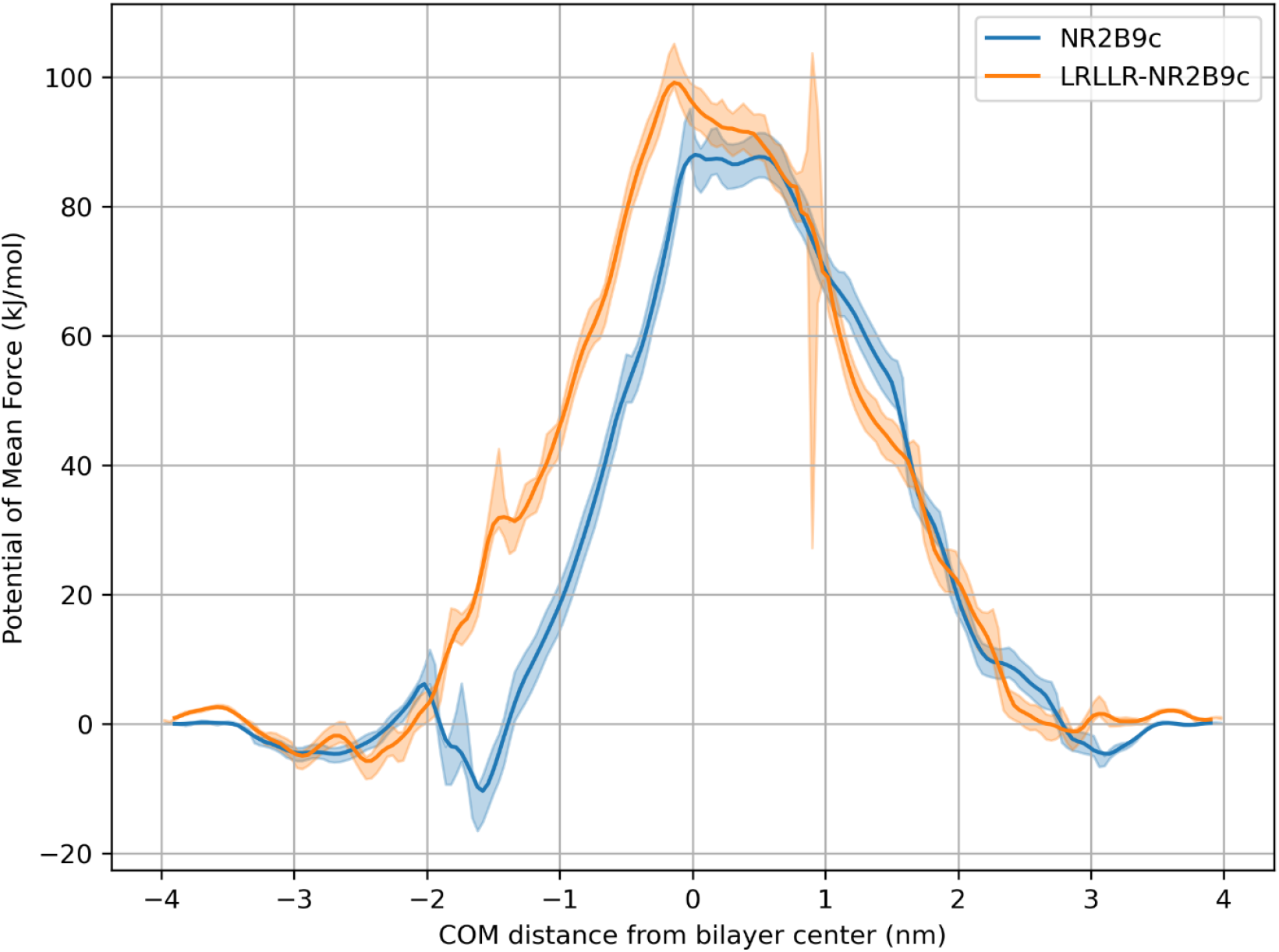
Potential of mean force profile for the NR2B9c peptide and its chimeric variation translocating a 90:10 POPC/POPG membrane. Shaded area around the profile represents a single standard deviation from the potential at that point.

**Figure 7:**
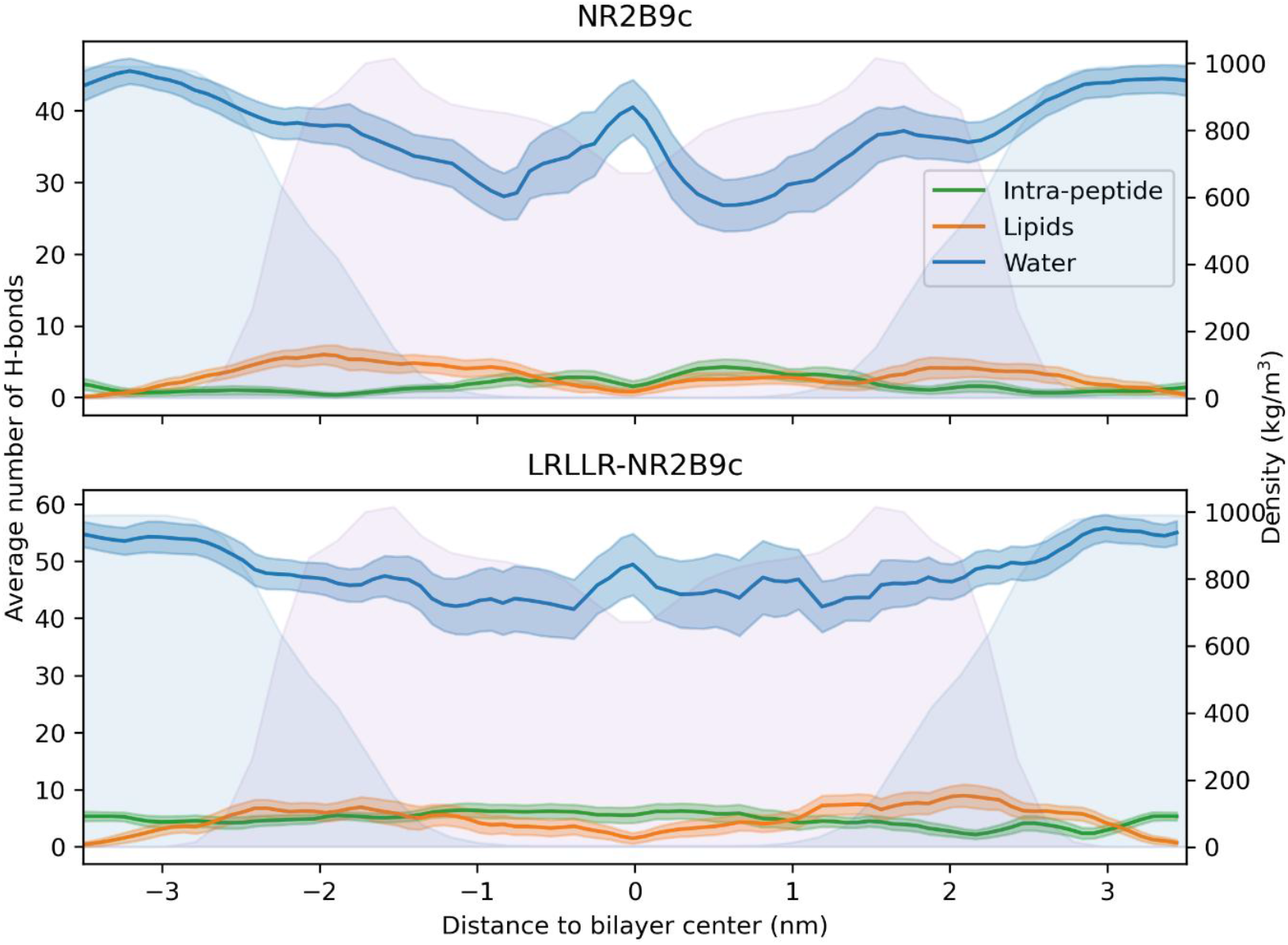
Number of hydrogen bonds involving the NR2B9c (top) and LRLLR-NR2B9c (bottom) peptides, separated by target. Shaded area in the background shows the water and lipid density as a function of the distance from the bilayer center.

Addition of the LRLLR motif to the C-terminus dramatically alters the energetic landscape of membrane interaction. The modification introduces a +2 charge and reduces overall hydrophobicity (GRAVY index: 1.31). Remarkably, the chimeric smacN-LRLLR peptide djisplays no translocation barrier; instead, the PMF reveals a deep energy well of −50 kJ/mol centered within the outer leaflet, approximately 1.2 nm from the bilayer center.

This stabilization arises from the ability of smacN-LRLLR to simultaneously satisfy both hydrophobic and electrostatic interactions. At the energy minimum, the added arginine residues maintain contact with negatively charged lipid headgroups while the leucine residues insert into the hydrophobic membrane core. The extended length of the chimeric peptide enables it to span sufficient distance to interact with both leaflets simultaneously when positioned at the bilayer center—a configuration inaccessible to the shorter unmodified peptide.

The modified peptide also exhibits enhanced structural order, adopting helical conformations 30% of the time compared to the complete absence of secondary structure in unmodified smacN. Neither peptide variant forms significant intramolecular hydrogen bonds during translocation. Notably, while smacN-LRLLR does not induce pore formation, a small number of water molecules co-translocate with the peptide.

For unmodified smacN positioned at the bilayer center, stabilization requires substantial membrane deformation: lipid headgroups must bend out of the bilayer plane to solvate the polar backbone, disrupting bilayer integrity and creating a water-permeable pore. This unfavorable membrane perturbation accounts for the significant energy barrier observed in the PMF.

### NR2B9c peptide

The NR2B9c nonapeptide (KLSSIESDV) carries a net negative charge (−1) at physiological pH and is slightly hydrophilic (GRAVY index: −0.09). This peptide functions by disrupting PSD-95 interactions through its C-terminus, which must remain accessible for target recognition. Addition of the LRLLR motif to the N-terminus reverses the net charge to +1 and shifts the hydropathy index to slightly hydrophobic (GRAVY index: 0.11).

Unmodified NR2B9c encounters a small barrier of +10 kJ/mol at the diffuse counterion layer before settling into a shallow interfacial well of −10 kJ/mol at 1.6 nm from the bilayer center. The translocation barrier reaches +85 kJ/mol. At the interfacial position, stability derives primarily from extensive hydration (40 ± 8 water hydrogen bonds) with only sporadic lipid interactions (4 ± 3 hydrogen bonds) and minimal intramolecular hydrogen bonding (<1). Residues Leu2, Ser4, and Ile5 insert partially into the hydrophobic region while maintaining peptide solvation.

Contrary to expectations, LRLLR-NR2B9c exhibits a *higher* translocation barrier of +100 kJ/mol. More strikingly, the chimeric peptide fails to achieve the interfacial stability observed for the unmodified sequence. At 1.6 nm from the bilayer center— where unmodified NR2B9c resides at its energy minimum—LRLLR-NR2B9c sits +30 kJ/mol above its bulk solution reference state.

Analysis of hydrogen bonding patterns reveals the structural basis for this destabilization. The LRLLR addition promotes intramolecular hydrogen bond formation (5 ± 1 bonds versus <1 for unmodified), which in turn favors helical structure (35% versus 20% helical content at the interface). This increased structural rigidity prevents optimal positioning of hydrophobic residues: the newly added leucines (particularly Leu1 and Leu4) cannot simultaneously insert into the membrane alongside the original hydrophobic residues (Leu7, Ser9, and Ile10, formerly Leu2, Ser4, and Ile5). Consequently, at least one hydrophobic side chain remains exposed to the aqueous environment, precluding stable interfacial association.

At the bilayer center, both peptide variants require pore formation for stabilization. In LRLLR-NR2B9c, the two arginine residues anchor to opposite leaflets (Arg5 to the outer leaflet, Arg2 to the inner), creating a stretched conformation. A bend at Ser8-Ser9 positions the polar Glu11 and Ser12 residues at the bilayer midplane, where insufficient polar contacts are available for stabilization, resulting in higher free energy compared to unmodified NR2B9c.

### Comparative analysis

The contrasting outcomes for smacN and NR2B9c chimeras illuminate the structural determinants of successful motif transfer. Three key factors emerge from comparison of these systems.

First, peptide length relative to bilayer thickness determines whether simultaneous interaction with both membrane interfaces is geometrically feasible. The smacN-LRLLR chimera (9 residues) achieves sufficient length to span the bilayer and engage both leaflets, with arginine residues contacting headgroups on one side while the N-terminus interacts with the opposite leaflet. The LRLLR-NR2B9c chimera (14 residues), despite greater length, adopts conformations that prevent equivalent bridging interactions.

Second, the structural propensity of the chimeric peptide critically influences translocation competence. The enhanced intramolecular hydrogen bonding and helical character induced in LRLLR-NR2B9c restricts conformational flexibility, preventing the hydrophobic residues from adopting optimal membrane-inserted orientations. In contrast, smacN-LRLLR maintains sufficient flexibility to accommodate favorable lipid interactions while gaining modest secondary structure that may aid in maintaining amphipathic character.

Third, the distribution of polar and charged residues along the chimeric sequence affects the energetics of bilayer positioning. In smacN-LRLLR, the hydrophobic smacN portion and the amphipathic LRLLR motif create a favorable gradient that allows progressive membrane insertion. In LRLLR-NR2B9c, the internal glutamate residue (Glu11) becomes positioned at the bilayer center during translocation, where it cannot be adequately stabilized, substantially raising the energetic cost.

These results demonstrate that successful transfer of cell-penetrating activity requires compatibility between the structural and physicochemical properties of the motif and the recipient peptide. Simple attachment of a penetrating motif does not guarantee enhanced translocation; indeed, as the NR2B9c case illustrates, such modification may prove counterproductive.

## Discussion

Our simulations demonstrate that the LRLLR motif, previously identified as essential for passive translocation of the TP1 cell-penetrating peptide, can confer membrane-translocating properties to other peptides—but only under specific structural and physicochemical conditions. Attachment of LRLLR to the C-terminus of smacN eliminated the translocation energy barrier entirely, replacing it with a favorable −50 kJ/mol energy well within the outer leaflet. In contrast, N-terminal LRLLR addition to NR2B9c increased the translocation barrier from +85 to +100 kJ/mol. These divergent outcomes reveal that successful motif transfer depends on the compatibility between the penetrating motif and the recipient peptide sequence.

### Structural Basis for smacN-LRLLR success

The successful enhancement of smacN membrane permeability by LRLLR addition can be attributed to favorable complementarity between the two sequence elements. The highly hydrophobic smacN tetrapeptide (GRAVY: 2.14) lacks any charged residues, presenting a surface that interacts favorably with the membrane core but provides no means of engaging with lipid headgroups or traversing the polar interfacial region. The LRLLR motif supplies precisely what smacN lacks: two positively charged arginine residues that anchor to anionic phosphate groups, interspersed with leucines that maintain hydrophobic continuity with the smacN portion.

The resulting chimera exhibits the hallmarks of an effective amphipathic peptide. When positioned at the energy minimum within the outer leaflet, smacN-LRLLR simultaneously satisfies hydrophobic contacts through its leucine-rich regions and electrostatic interactions through its arginine residues. The peptide length proves sufficient to bridge both leaflets during transit through the bilayer center, avoiding the need for membrane deformation or pore formation that would impose additional energetic costs.

The modest increase in helical propensity (from absent to 30%) provides structural benefit without imposing excessive rigidity. This secondary structure likely helps to maintain the spatial segregation of polar and nonpolar residues characteristic of amphipathic helices, while retaining sufficient flexibility to adapt to the varying polar environment across the membrane depth.

The C-terminal placement of LRLLR leaves the N-terminus of smacN—the recognition element for inhibitor of apoptosis proteins—unobstructed and available for target engagement upon cellular entry. This design consideration proves fortuitously compatible with the translocation mechanism, as our simulations reveal that the free N-terminus participates in the final stage of translocation by engaging with inner leaflet phosphates.

### Structural Basis for NR2B9c-LRLLR failure

The failure of LRLLR to enhance NR2B9c translocation stems from multiple incompatibilities between the motif and its recipient. Unlike the short, uniformly hydrophobic smacN, NR2B9c presents a more complex sequence containing internal polar (Ser3, Ser4, Ser7, Ser8) and charged (Lys1, Glu6, Asp8) residues. These existing polar elements preclude the clean amphipathic organization that LRLLR achieves with smacN.

Most critically, the addition of LRLLR promotes intramolecular hydrogen bonding and helical structure in the chimeric peptide. While secondary structure generally favors membrane interaction for amphipathic sequences, in this case the increased rigidity prevents optimal positioning of hydrophobic residues. The original hydrophobic anchors of NR2B9c (Leu2, Ile5) cannot simultaneously insert into the membrane alongside the newly introduced leucines of the LRLLR motif. This geometric incompatibility leaves hydrophobic surface area exposed to water at the interface—an energetically unfavorable configuration that destabilizes the chimera relative to unmodified NR2B9c.

The internal glutamate residue (Glu6 in NR2B9c, Glu11 in the chimera) presents an additional complication. During translocation, the extended conformation adopted by LRLLR-NR2B9c—with arginines anchored to opposite leaflets—positions this negatively charged residue at the bilayer midplane, where polar stabilization is minimal. The energetic penalty for burying this charge contributes substantially to the elevated translocation barrier.

These findings suggest that TAT-NR2B9c, the conjugate currently in clinical development, may represent a more suitable approach for this particular therapeutic peptide. The TAT peptide, with its extended length and multiple arginine residues, likely employs different translocation mechanisms—potentially including membrane pore formation or recruitment to endocytic pathways—that circumvent the interfacial stability problems we observe with LRLLR addition.

### General principles for motif transfer

Our comparative analysis suggests several principles that may guide future efforts to transfer cell-penetrating activity to therapeutic peptides:

#### Sequence complementarity

The penetrating motif should supply functional groups absent in the recipient peptide. LRLLR provides both positive charges (arginine) and hydrophobic bulk (leucine). Recipients lacking either element may benefit; recipients possessing both may gain less advantage.

#### Conformational compatibility

The chimeric peptide must retain sufficient flexibility to adopt membrane-favorable conformations. If motif addition induces stable secondary structures that constrain hydrophobic residue positioning, translocation may be impaired rather than enhanced.

#### Polar residue distribution

Internal charged or highly polar residues within the recipient sequence may become trapped in unfavorable positions during translocation. Short, uniformly hydrophobic recipients present lower risk of such complications.

#### Length considerations

The chimeric peptide should achieve sufficient length to span the bilayer when extended, enabling simultaneous engagement with both leaflets. However, excessive length may introduce conformational complexity that undermines this advantage.

#### Attachment site selection

The motif attachment site should preserve the functional epitope of the therapeutic peptide while placing the penetrating elements where they can engage the membrane effectively. In the successful smacN-LRLLR case, C-terminal attachment left the N-terminal recognition sequence intact while allowing the LRLLR arginines to interact with headgroups during entry.

### Comparison to related studies

The translocation barriers obtained in our simulations are consistent with values reported in the literature for other cell-penetrating peptides. Huang and García reported barriers of 80–100 kJ/mol for oligoarginine translocation across DPPC bilayers, comparable to our observations for LRLLR and unmodified NR2B9c^54^. Similarly, umbrella sampling studies by Yesylevskyy et al. found interfacial energy wells of 20–60 kJ/mol for various CPPs^55^, consistent with our values for LRLLR (−40 kJ/mol) and smacN-LRLLR (−50 kJ/mol).

The −50 kJ/mol energy well observed for smacN-LRLLR is notably deep compared to most reported CPP-membrane interactions, suggesting that this chimera may exhibit exceptionally stable membrane association. Whether this represents an advantage (enhanced membrane accumulation preceding translocation) or a limitation (kinetic trapping at the interface) remains to be determined experimentally.

Our observation that LRLLR requires pore formation during the transition state aligns with the work of Herce and García, who demonstrated water pore involvement in TAT translocation^56^. However, the relatively modest number of water molecules (20 ± 5) involved in LRLLR translocation suggests a smaller pore compared to the larger penetrations reported for arginine-rich CPPs, potentially reflecting the lower charge density of the LRLLR motif.

The translocation barriers we calculate (60–100 kJ/mol) exceed values compatible with spontaneous passive translocation on biological timescales. This apparent discrepancy with experimental observations of LRLLR translocation likely reflects the absence of membrane potential in our simulations. Several theoretical and computational studies have demonstrated that physiological membrane potentials can substantially reduce translocation barriers for cationic peptides by 20–40 kJ/mol [ref, ref], which would bring our calculated barriers into better agreement with experimental translocation rates.

Additionally, our results do not account for the potential role of megapolarization events near ion channels, which have been proposed to facilitate CPP entry through transient local electric field amplification^57^. If LRLLR translocation depends on such events in vivo, the barriers we calculate would represent upper bounds rather than operative values.

### Future directions

These computational findings generate several experimentally testable predictions. The most direct test would compare the membrane permeability of smacN versus smacN-LRLLR using fluorescently labeled variants in liposome leakage assays or cell uptake experiments. Our simulations predict substantially enhanced uptake for the chimeric peptide, with preferential accumulation within the membrane phase.

Parallel experiments with LRLLR-NR2B9c should demonstrate no enhancement—or possibly reduced uptake—compared to unmodified NR2B9c. Surface plasmon resonance or isothermal titration calorimetry could test our prediction that LRLLR-NR2B9c exhibits weaker membrane affinity than the unmodified peptide.

Circular dichroism spectroscopy in membrane-mimetic environments could validate our predictions regarding secondary structure differences between successful and unsuccessful chimeras. We predict higher helical content for smacN-LRLLR compared to smacN, and we predict that LRLLR-NR2B9c secondary structure may correlate with reduced membrane affinity.

From a computational perspective, several extensions of this work merit consideration. Testing motif transfer across different membrane compositions— including higher anionic lipid content, cholesterol inclusion, and asymmetric leaflet compositions—would evaluate the robustness of our conclusions. The influence of membrane potential could be explicitly modeled using charge imbalance methods or applied electric field approaches, potentially reconciling our calculated barriers with experimental translocation rates.

The introduction of flexible spacers between the therapeutic peptide and the penetrating motif represents a promising avenue for optimization. A short glycine-rich or aminohexanoic acid linker might allow LRLLR to function more independently of the recipient sequence, potentially rescuing the failed NR2B9c case by reducing conformational coupling between the two elements.

Finally, systematic evaluation of motif transfer across a library of therapeutic peptides would enable identification of sequence features predictive of success or failure. Such a dataset could support machine learning approaches to predict motif transfer compatibility without requiring exhaustive simulation campaigns, substantially accelerating the design cycle for cell-penetrating peptide conjugates.

## Conclusions

This study establishes that cell-penetrating motif transfer is feasible but sequence-dependent. The LRLLR motif alone exhibits limited spontaneous translocation capacity; despite demonstrating strong interfacial affinity (−40 kJ/mol), the isolated pentapeptide faces a substantial barrier (+100 kJ/mol from the interfacial minimum) that likely requires additional factors—such as membrane potential, cargo interactions, or megapolarization events—for spontaneous crossing.

The contrasting outcomes observed for the two therapeutic peptides reveal that compatibility between motif and recipient sequence determines success. Addition of LRLLR to the C-terminus of smacN transforms an unfavorable energetic profile (+65 kJ/mol barrier) into a highly favorable one (−50 kJ/mol energy well within the outer leaflet), representing a dramatic enhancement of membrane permeability. In contrast, N-terminal LRLLR addition to NR2B9c increases rather than decreases the translocation barrier (from +85 to +100 kJ/mol) and destabilizes interfacial binding. These divergent results demonstrate that successful transfer requires complementarity in charge distribution, hydrophobicity, and conformational flexibility. Short, neutral, hydrophobic recipients such as smacN represent favorable platforms, whereas longer peptides with internal polar residues and propensity for rigid secondary structures may prove incompatible.

The smacN-LRLLR chimera warrants experimental validation based on these findings. The complete elimination of the translocation barrier, combined with preservation of the N-terminal recognition motif required for inhibitor of apoptosis protein targeting, positions this conjugate as a promising candidate for chemotherapy-resistant cancers where IAP overexpression contributes to treatment failure. Experimental characterization of membrane permeability, cellular uptake, and target engagement is now justified.

Finally, the divergent outcomes observed for smacN and NR2B9c demonstrate the value of computational screening in CPP conjugate design. Umbrella sampling simulations can distinguish promising candidates from likely failures prior to experimental investment, and the negative result for NR2B9c holds importance beyond the specific system studied—it illustrates that simple attachment of cell-penetrating sequences may prove counterproductive for certain cargo peptides. Integration of such predictive methods into early-stage design pipelines may substantially improve success rates and accelerate development of membrane-permeable therapeutic peptides.

## Acknowledgements

D. Munoz-Gacitua acknowledges support from a postdoctoral fellowship (POSTDOC_DICYT 022443BA) granted by the Vicerrectoría de Investigación, Innovación y Creación of Universidad de Santiago de Chile (USACH). The authors acknowledge and thank Fundación Biociencia for providing the computational resources that made this research possible.

## Conflict of interest statement

The authors declare no conflicts of interest. This study was fully funded by Fundación Biociencia and Universidad de Santiago de Chile (USACH). The funders had no role in the design of the study; in the collection, analyses, or interpretation of data; in the writing of the manuscript; or in the decision to publish the results.

## Credit attribution statement

D. Munoz-Gacitua conceived the research, designed the methodology, developed the tools for simulations and posterior analysis, and drafted the original manuscript. J. Blamey provided essential research resources, supervised the project, and critically reviewed and edited the manuscript. All authors approved the final version for publication.

## AI assistance statement

During the preparation of this manuscript, the authors utilized a locally self-hosted large language model (LLM) to enhance grammar, sentence structure, and readability. To ensure data governance and maintain confidentiality of unpublished research, all AI assistance was performed using Biociencia’s self-hosted LLM infrastructure, preventing any external data transmission. All scientific content, methodology, data analysis, interpretations, and conclusions remain the original intellectual work of the authors, who take full responsibility for the accuracy and integrity of this publication.

## Notes

### Competing Interest Statement

The authors have declared no competing interest.

### Summary of Updates

docs: Adds supplementary information on simulation time per peptide, and sampling of each translocation study performed

