## Supplementary information for "Sequence-dependent transferability of the LRLLR membrane translocation motif: A computational study of smacN and NR2B9c peptides"

### Supplementary material

Table S1: Total simulation time per peptide

| *Peptide* | *Transient simulation time [μs]* | *Production simulation time [μs]* |
|---|---|---|
| *LRLLR* | 2.14 | 3.92 |
| *NR2B9c* | 3.94 | 5.69 |
| *LRLLR-NR2B9c* | 2.05 | 3.40 |
| *smacN* | 2.06 | 4.33 |
| *smacN-LRLLR* | 3.28 | 6.34 |
| **Total** | **13.47** | **23.68** |

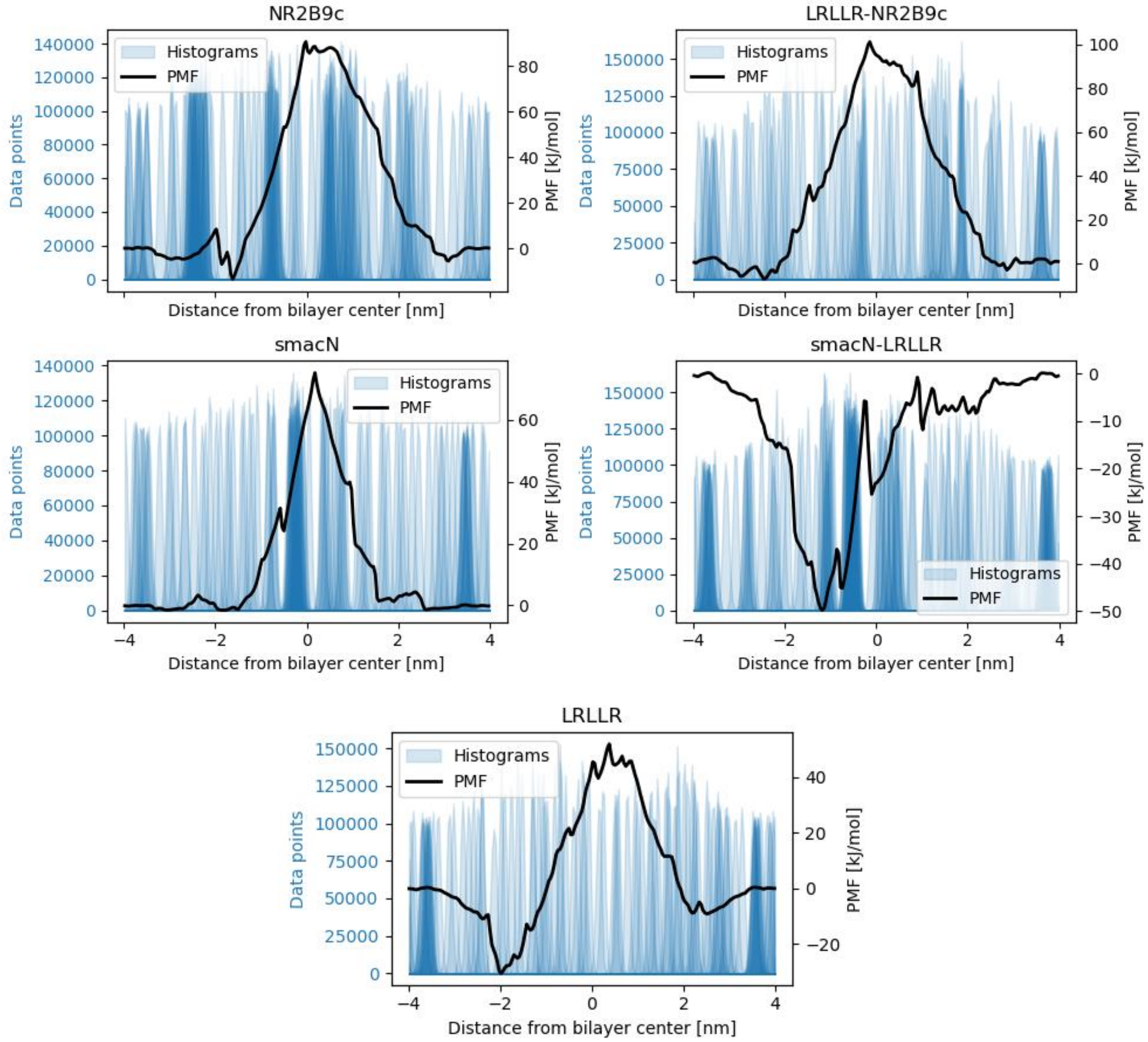


Figure S2: Potential of mean force plots for the process of peptides translocating a membrane as calculated from an Umbrella Sampling/WHAM procedure. Shown in blue, the histograms detailing the sampling at each distance.
